# Category trumps shape as an organizational principle of object space in the human occipitotemporal cortex

**DOI:** 10.1101/2022.10.19.512675

**Authors:** Elahe’ Yargholi, Hans Op de Beeck

## Abstract

The organizational principles of the object space represented in human ventral visual cortex are debated. Here we contrast two prominent proposals that, in addition to an organization in terms of animacy, propose either a representation related to aspect ratio or to the distinction between faces and bodies. We designed a critical test that dissociates the latter two categories from aspect ratio and investigated responses from human fMRI and deep neural networks (BigBiGAN). Representational similarity and decoding analyses showed that the object space in occipitotemporal cortex (OTC) and BigBiGAN was partially explained by animacy but not by aspect ratio. Data-driven approaches showed clusters for face and body stimuli and animate-inanimate separation in the representational space of OTC and BigBiGAN, but no arrangement related to aspect ratio. In sum, the findings go in favor of a model in terms of an animacy representation combined with strong selectivity for faces and bodies.

## Introduction

Visual object recognition is crucial for human image understanding. Neuroscientists have shown that the lateral and ventral occipitotemporal cortex (OTC) is particularly important for object recognition. This large cortical region is a primary example of the complexity of functional selectivity in the human brain. Studies have uncovered large-scale maps for distinctions such as animate versus inanimate, selective areas for specific categories like faces, bodies, and scenes, and further maps for visual and semantic features^1–5^. However, it is unclear how these different aspects of this functional architecture can be put together in one comprehensive model.

Over the years, there have been very diverse proposals about the principles governing object space in OTC^6^. For example, some studies suggest that low-level visual properties could explain the functional organization of OTC for object recognition^7,8^, while others identified a central role for higher-level features, particularly in more anterior regions^2,9–13^. Overall, studies support a rich representation of objects that combines selectivity for features from multiple levels of complexity^14^.

Recently, ref.^15^ proposed a comprehensive map of object space in monkeys’ inferior temporal (IT) cortex based upon data from monkey fMRI, single-neuron electrophysiology, and deep neural network (DNN). After identifying a network of object-selective areas beyond the face- and body-selectivity areas through fMRI, they sampled single neurons distributed across areas with different population-level selectivity and suggested that the IT cortex is organized as a map with two main dimensions, aspect ratio (stubby-spiky) and animate-inanimate. This map would explain the location of specific regions in the IT cortex. In particular, the anatomical location of face- and body-selective regions would be constrained by the visual properties of these stimuli, with faces being stubby and bodies being spiky. In addition, ref.^15^ found that individual IT cells map stimuli to the two axes of this space (stubby-spiky and animate-inanimate) and claimed a unified picture of the IT organization. A similar space was observed in DNNs, further confirming earlier findings of correspondences between primate vision and DNNs^16–21^.

This proposal of a two-dimensional animacy x aspect ratio space is appealing as a comprehensive model. At first sight, it fits with earlier but somewhat dispersed findings of both animacy and spiky-stubby selectivity in human and monkey^11,14,22^. In addition, the two-dimensional object space as proposed by ref.^15^ was mostly confirmed in the human brain with the same stimulus set^23^. Nevertheless, the evidence for the model is fundamentally flawed. None of the experiments properly controlled the two dimensions. In ref.^15^ the stimulus set overall dissociated stubby-spiky from animate-inanimate, but this was not true within stimulus classes: Face stimuli were all stubby and body stimuli were mostly spiky. Much of the evidence for aspect ratio as an overall dimension and as a dimension underlying face and body selectivity and determining the location of selectivity for these categories, might be due to this major confound between aspect ratio and stimulus category (face versus body). It is possible that, alternatively, face versus body selectivity has a primary role in the organization of OTC and that aspect ratio is at best a dimension of secondary importance. This is an alternative view of how object representations are organized, why they have this structure, and what constraints might determine where specific category-selective regions emerge in OTC.

We designed a new experimental paradigm in order to dissociate these two alternative hypotheses: An object space organized in terms of the dimensions of animacy and aspect ratio, or instead an object space organized in terms of a primary dimension of animacy and a further distinction between faces and bodies that is NOT related to the aspect ratio of these categories. To be able to properly distinguish among these alternatives, we designed a novel stimulus set in which the face and body categories are explicitly dissociated from aspect ratio (being stubby or spiky). We obtained neural responses with this paradigm through functional magnetic resonance imaging (fMRI), and examined object representations in human OTC as well as a state-of-the-art DNN. We observed no tuning representation for aspect ratio in human face and body regions. A representation for aspect ratio was restricted to right object-selective regions and only present for inanimate objects. Aspect ratio failed convincingly as a candidate for the status of encompassing dimension of object space, and this both in humans and DNNs. In sum, the findings go against the two-dimensional animacy x aspect ratio model, and in favor of a model in terms of an animacy representation combined with strong selectivity for faces and bodies.

## Results

In this study, we aimed to investigate if animacy and aspect ratio are the two main dimensions organizing object space in OTC using a stimulus set that properly dissociated the type of visual stimuli, and in particular face and body, from aspect ratio. A stimulus set of body, face, manmade and natural objects with a comparably wide range of aspect ratios in each of these four categories was presented (Fig. 1A), and fMRI responses (beta weights) were extracted in each voxel of the brain for each stimulus. Then, we employed fMRI multivariate (representational similarity analysis and classification) and univariate analysis. Representations of our stimuli in a DNN’s latent space were also inspected for the animacy and aspect ratio contents.

**Figure 1.**
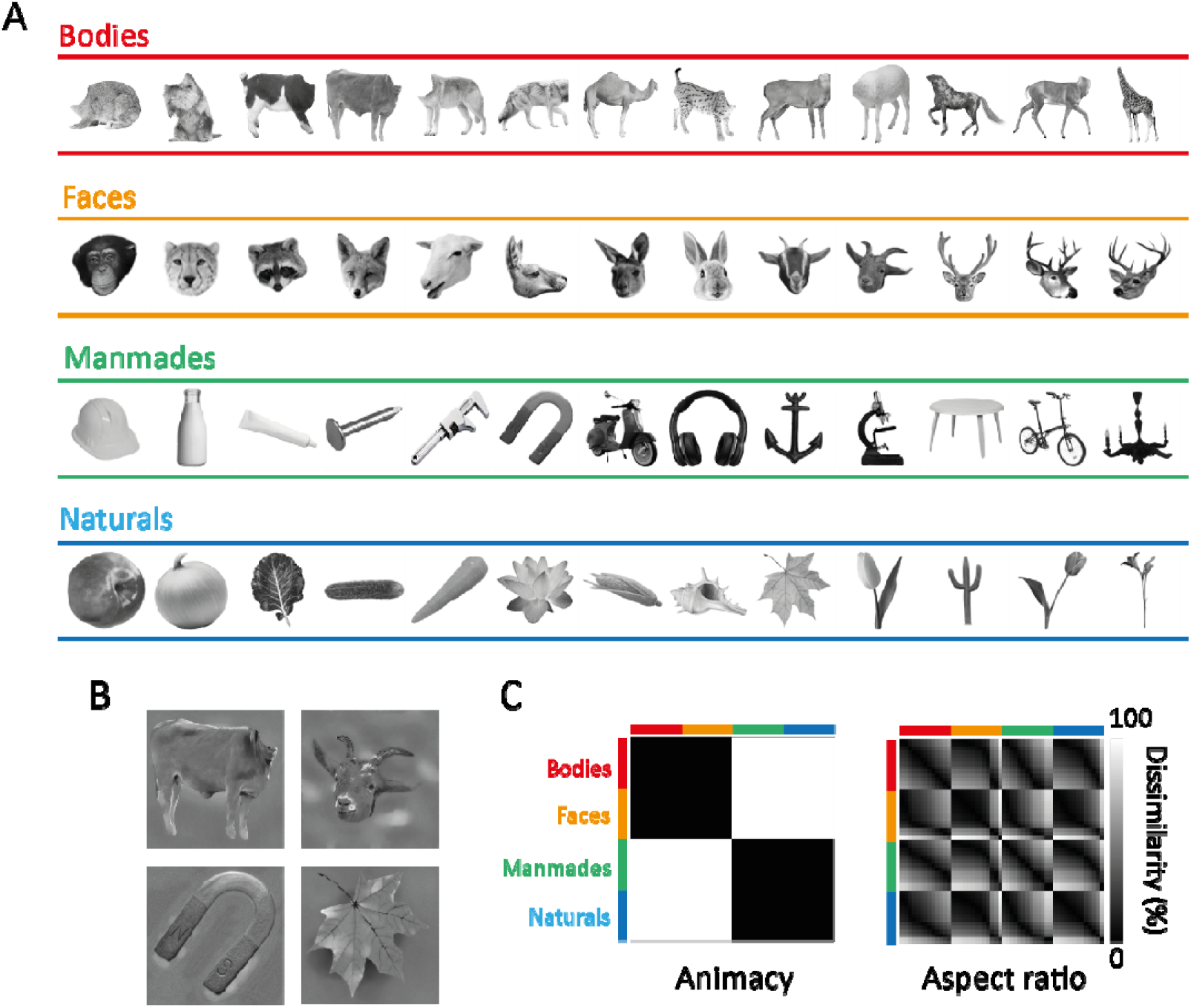
Stimulus design. **A**. All 52 images used to prepare stimuli, in the order of increasing aspect ratio from left to right and color-coded based on the category. **B**. Examples of finalized stimuli (gray background, equalized luminance histogram, etc.) used for fMRI experiment and computational analysis. **C**. The two main model RDMs used throughout the research. The axes of RDMs are color-coded based on stimulus category.]

### To what extent do animacy and aspect ratio models explain the organization of OTC’s object space?

To examine the neural representation of animacy and aspect ratio in different ROIs, we compared neural dissimilarity in the multivoxel patterns elicited by individual images with predictions from animacy and aspect ratio models (Fig 1C). The result for OTC (combination of body, face, and object selective ROIs) is shown in Fig. 2A. Both left and right OTC showed significant positive correlations with the animacy model (one-sided one sample t-test, left OTC: t(14) = 10.39, p < 0.001, right OTC: t(14) = 13.09, p < 0.001) and not with the aspect ratio model (one-sided one sample t-test, left OTC: t(14) = 0.0054, p > 0.5, right OTC: t(14) = −0.38, p > 0.5). The direct comparison of the correspondence with neural dissimilarity between animacy and aspect ratio showed significant differences in both hemispheres (two-sided paired t-test, left OTC: t(14) = 10.02, p < 0.001, right OTC: t(14) = 11.38, p < 0.001), revealing a far stronger correspondence for the animacy model than for the aspect ratio model. The large effect size of this difference is further illustrated by the observation that all 15 participants showed a larger correspondence with animacy compared to the aspect ratio, and this in each hemisphere. The MDS plots for left and right OTC (Fig. 2B) reveal clusters for faces and bodies and a clear separation of animates from inanimates, but spiky and stubby stimuli are intermingled and not at all separated in clusters.

**Figure 2.**
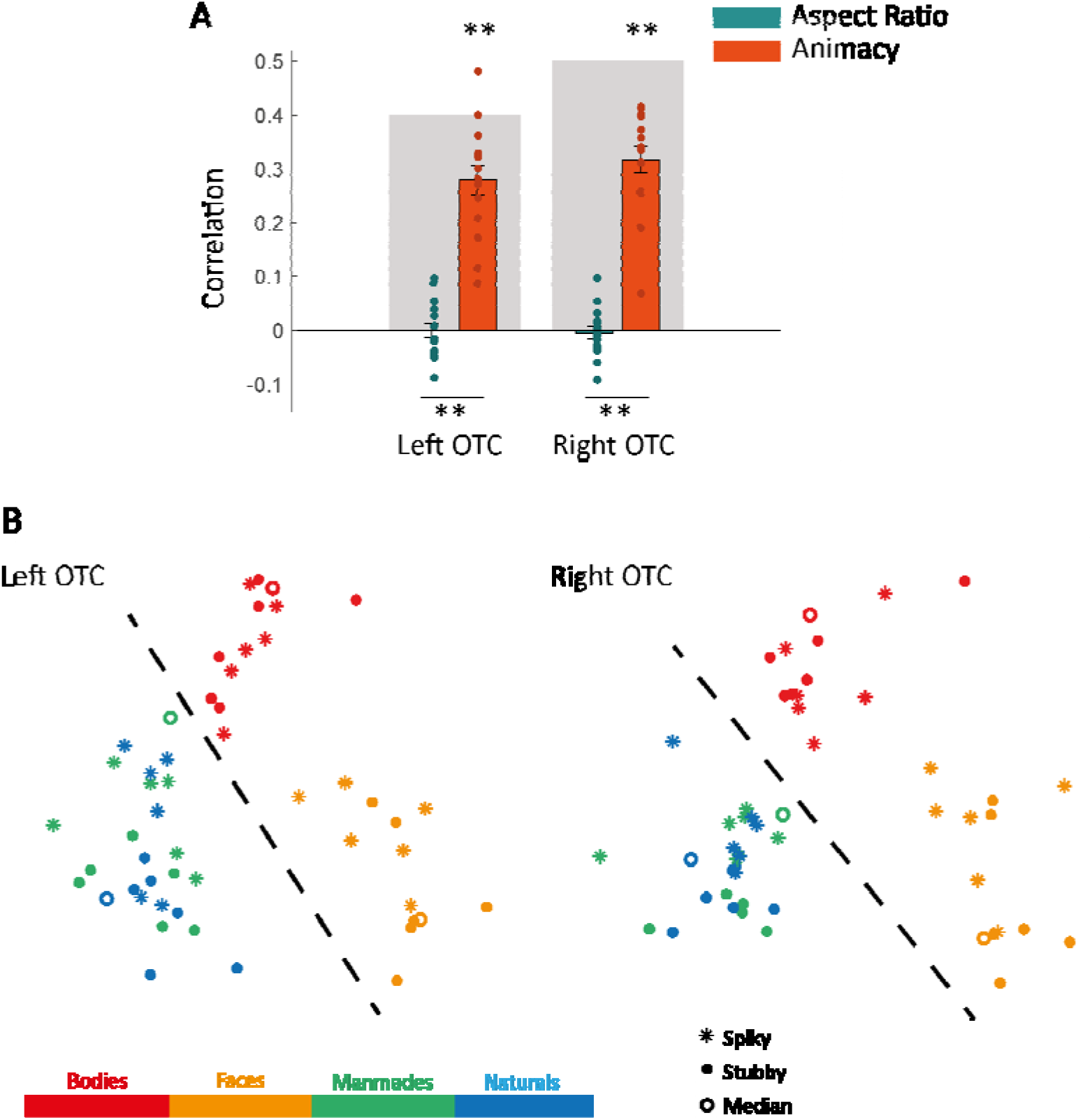
Representational similarity and effects of aspect ratio and animacy in the occipitotemporal cortex. **A**. The bar plot shows the mean Spearman’s correlations between neural RDMs for individual subjects (dots) and model RDMs. Error bars indicate standard errors of the means and gray background bars indicate the noise ceiling. One-sided one sample or two-sided paired t-test, ** p < 0.001. **B**. Two-dimensional representational space as obtained by applying multidimensional scaling to the OTC’s neural RDM. Points are color-coded based on stimulus category and are dots, rings, or asterisks based on aspect ratio. The dashed line shows the separation of animates from inanimates.]

We followed the same approach for category-selective regions (Fig. 3) and the result showed significant correlations between neural dissimilarity and the animacy model in most ROIs (Fig. 4, one-sided one sample t-test, FDR-corrected across 26 ROIs, *: t > 4.05, p < 0.01, **: t > 5.31, p < 0.001), while no ROI had a significant correlation with aspect ratio. The direct comparisons of correlations with the two models also showed significant differences in most ROIs (Fig. 4, two-sided paired t-test, FDR-corrected across 26 ROIs, *: t > 4.32, p < 0.01, **: t > 5.22, p < 0.001).

**Figure 3.**
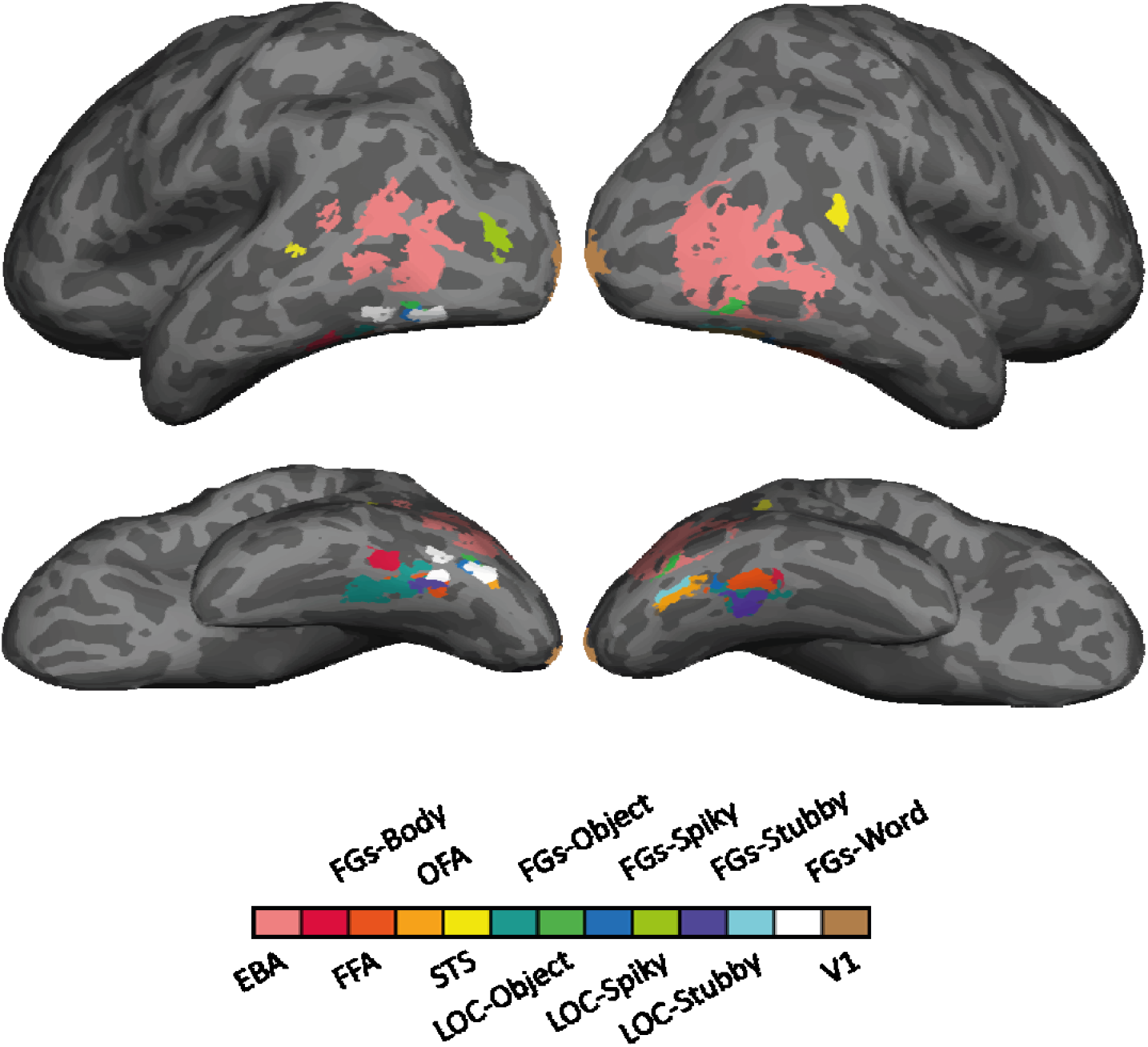
Inflated brain surface from an individual participant illustrating the category-selective ROIs examined, in lateral and ventral views.]

**Figure 4.**
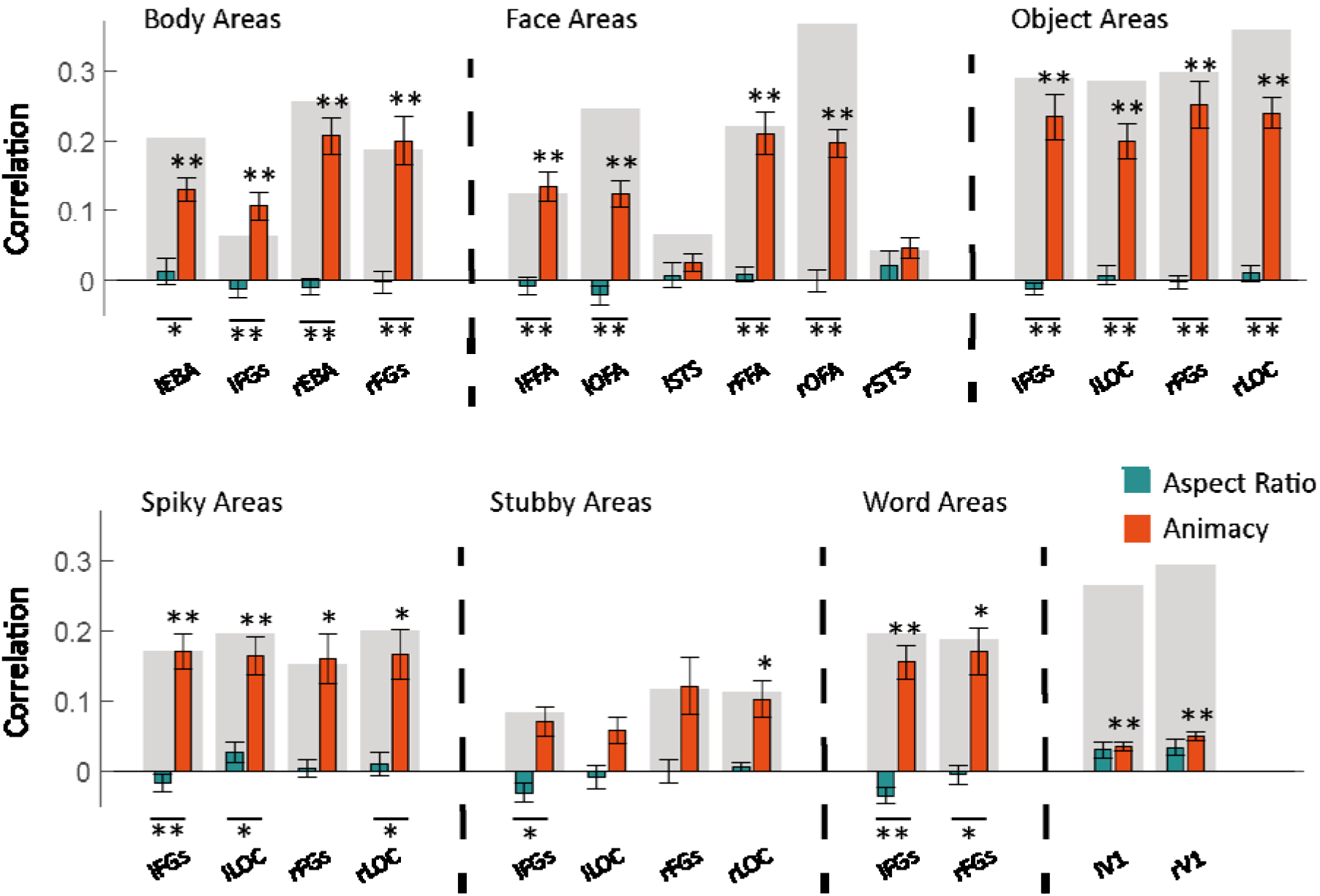
Effect of aspect ratio and animacy in specific category-selective ROIs. The bar plots show the mean Spearman’s correlations between neural RDMs for individual subjects and model RDMs. Error bars indicate standard errors of the means and gray background bars indicate the noise ceiling. One-sided one sample or two-sided paired t-test, FDR-corrected, ** p < 0.001, * p<0.01.]

The total absence of predictive power of the aspect ratio model is surprising and was scrutinized further. We hypothesized that it might exist only for the preferred category of a region of interest. For body, face, and object selective ROIs, we examined the correlation between neural dissimilarity and the aspect ratio model limiting stimuli to their favorite/selected category. Result of this analysis showed an effect of aspect ratio that was restricted to the right rFGs-object and rLOC-object (Fig. 5, one-sided one sample t-test, FDR-corrected across 4 ROIs, rFGs-object: t(14) = 2.97, p =0.0212, right OTC: t(14) = 3.18, p =0.0212). Looking more closely at objects (manmade and natural) in the MDS plot for the right OTC (Fig. 2B), we could also see a modest degree of arrangement in spiky and stubby stimuli. Thus, in accordance with previous work on shape representations with artificial shapes (e.g. ref.^22^), the aspect ratio of inanimate objects is represented in object-selective cortex. However, as the main results above show, aspect ratio is not an encompassing dimension when we also include animate stimuli (faces & bodies), and is thus unlikely to be an explanation for the location of face- and body-selective regions.

**Figure 5.**
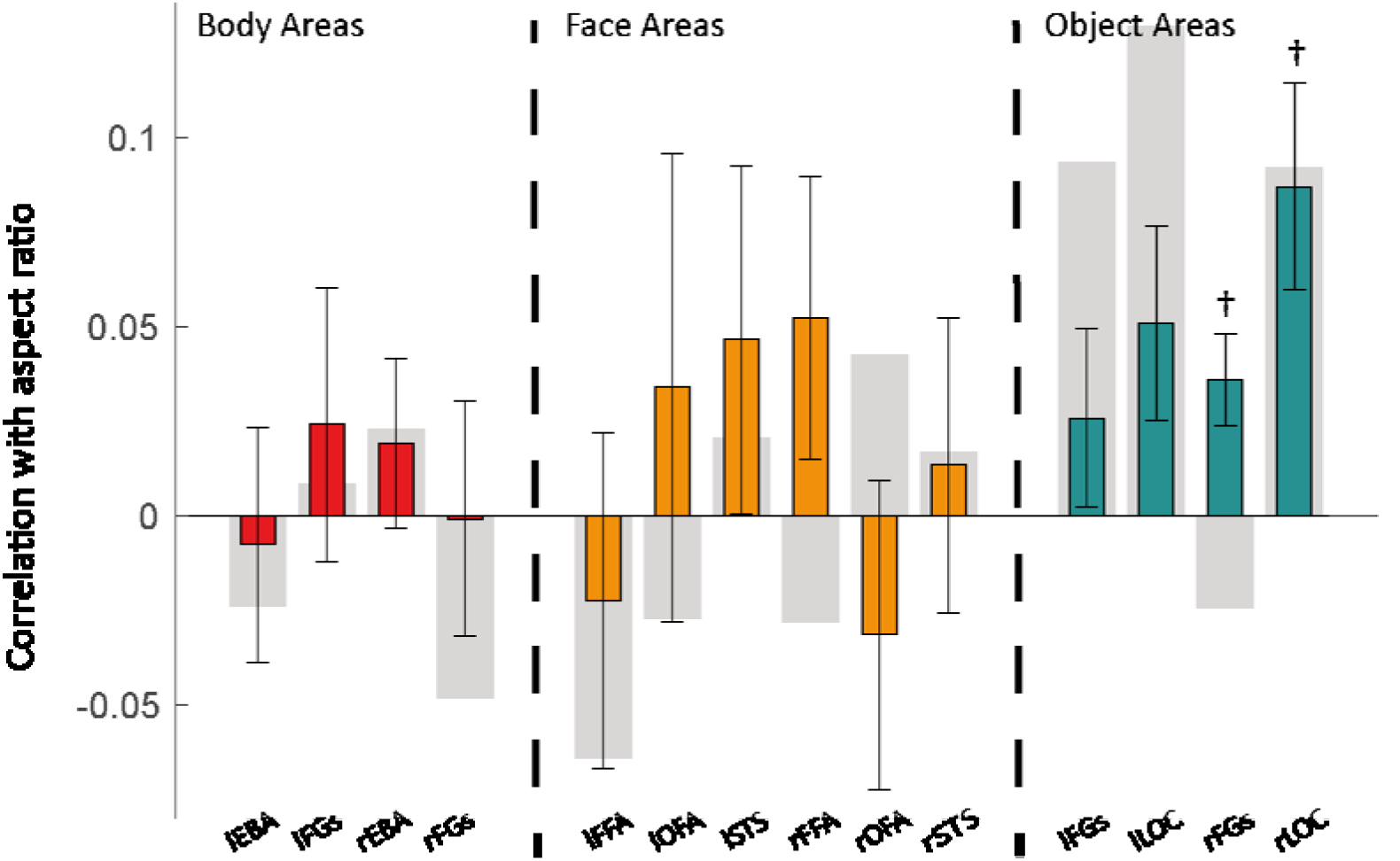
Effect of aspect ratio for the preferred stimulus category in body, face, and object selective ROIs. The bar plot shows the mean Spearman’s correlations between neural RDMs for individual subjects and the aspect ratio model RDM. Error bars indicate standard errors of the means and gra background bars indicate the noise ceiling. One-sided one sample t-test, FDR-corrected, † p<0.05.]

We also considered that the relative role of aspect ratio might shift along the ventral processing pathway. To explore this possibility, we investigated the similarity of neural and model dissimilarity in small consecutive ROIs along the ventral visual cortex (Fig. 6). As we move from posterior to anterior ROIs in both hemispheres, correlations between neural and animacy RDMs increased to significant levels and then decreased (Fig. 6, one-sided one sample t-test, FDR-corrected across 74 ROIs, thin line: t(14) > 3.78, p < 0.01, thick line: t(14) > 5.19, p < 0.001). In contrast, we did not observe significant correlations between neural and aspect ratio RDMs. Comparing the correlations with the two models also showed a significancy pattern similar to that of correlations between neural and animacy RDMs.

**Figure 6.**
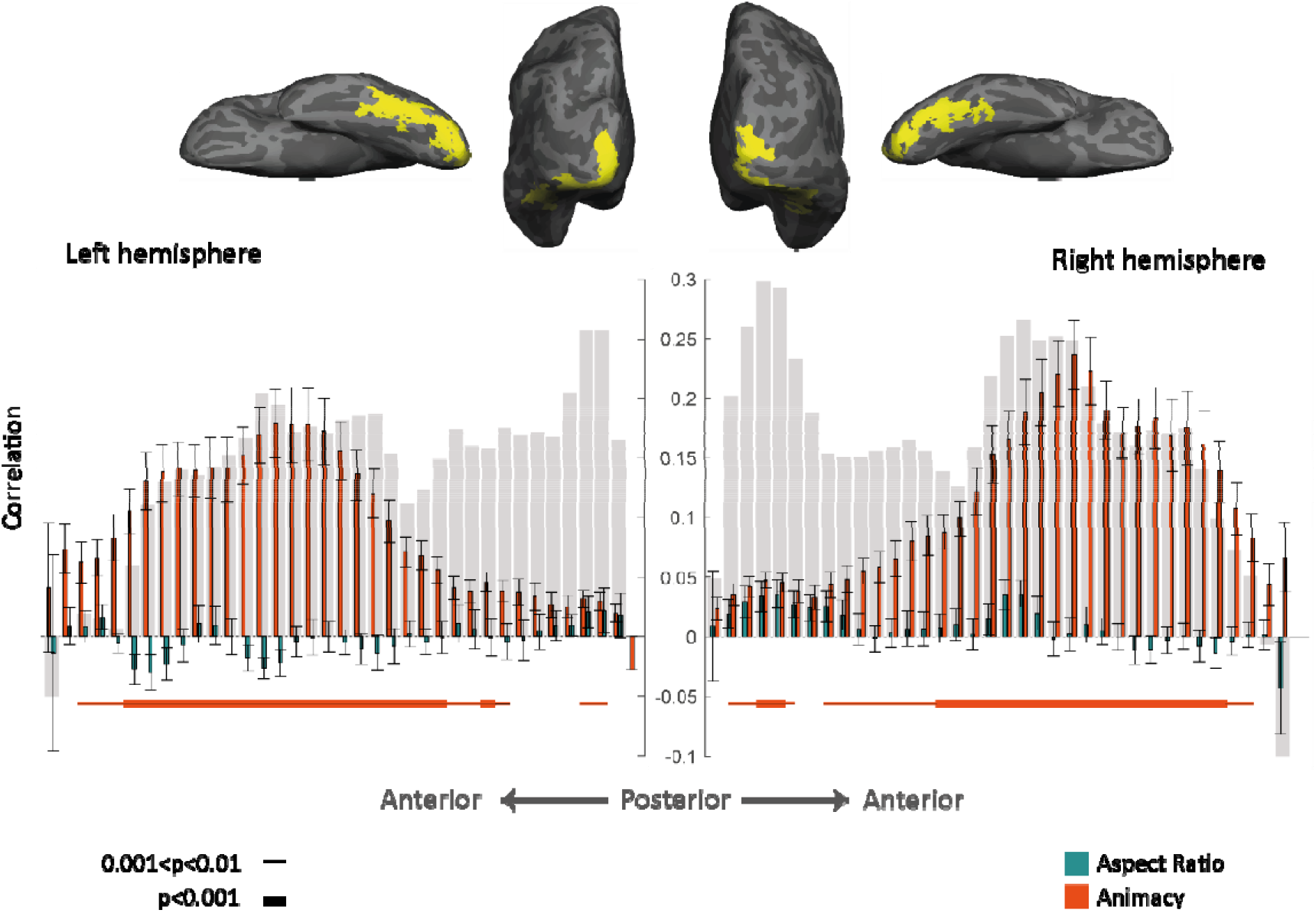
Gradual progression of effects of animacy and aspect ratio along the posterior-anterior gradient in ventral visual cortex. (top) Inflated brain surface from a representative participant showing the ventral visual cortex (union of small consecutive ROIs) in posterior and ventral views. (bottom) Comparing the model RDMs to the neural RDM of ROIs along the ventral visual cortex. The bar plot shows the mean Spearman’s correlations between neural RDMs for individual subjects and model RDMs (for the left hemisphere, the bar plot is mirrored to have a consistent direction with the anatomical image at the top). Error bars indicate standard errors of the means and gray background bars indicate the noise ceiling. One-sided one sample t-test, FDR-corrected, thin line p < 0.01, thick line p < 0.001]

In sum, these results suggest that object space in the whole OTC and most of OTC’s category-selective ROIs was much better explained by the animacy rather than the aspect ratio model and restricting stimuli to selected categories does not change this correlation pattern. Furthermore, this animacy content increases and then decreases as we move along the anatomical posterior-to-anterior axis in VOTC. All analyses confirm a major role for animacy as a primary dimension that determines the functional organization of object space. In contrast, the aspect ratio doesn’t function as a fundamental dimension characterizing the full extent of object space and its anatomical organization.

### Animacy distinction generalizes over aspect ratio, but not vice versa

In order to further investigate how each dimension is represented relative to the other, we examined the capability of neural representations to generalize distinctions along one dimension over the other dimension. With this aim, we compared within and cross-category classification accuracies for animate vs. inanimate and spiky vs. stubby dimensions. For animate vs. inanimate classification-within aspect ratio, we trained and tested classifiers with either spiky or stubby stimuli while for cross aspect ratio, classifiers were trained with spiky and tested with stubby stimuli and vice versa (supplemental table S2). For spiky vs. stubby classification-within animacy, we trained and tested classifiers with either animate or inanimate stimuli while for cross animacy, classifiers were trained with animate and tested with inanimate stimuli and vice versa (supplemental table S2).

As fig. 7 shows most category-selective ROIs had classification accuracies significantly above chance level for animate vs. inanimate-within and cross aspect ratio (Fig 7, one-sided one sample t-test, FDR-corrected across 26 ROIs, *: t > 3.9, p < 0.01, **: t > 4.98, p < 0.001). In contrast, only a few ROIs had classification accuracies significantly above the chance level for spiky vs. stubby-within animacy and they were not able to generalize the classification across animacy (Fig 8, one-sided one sample t-test, FDR-corrected across 26 ROIs, *: t > 3.9, p < 0.01).

**Figure 7.**
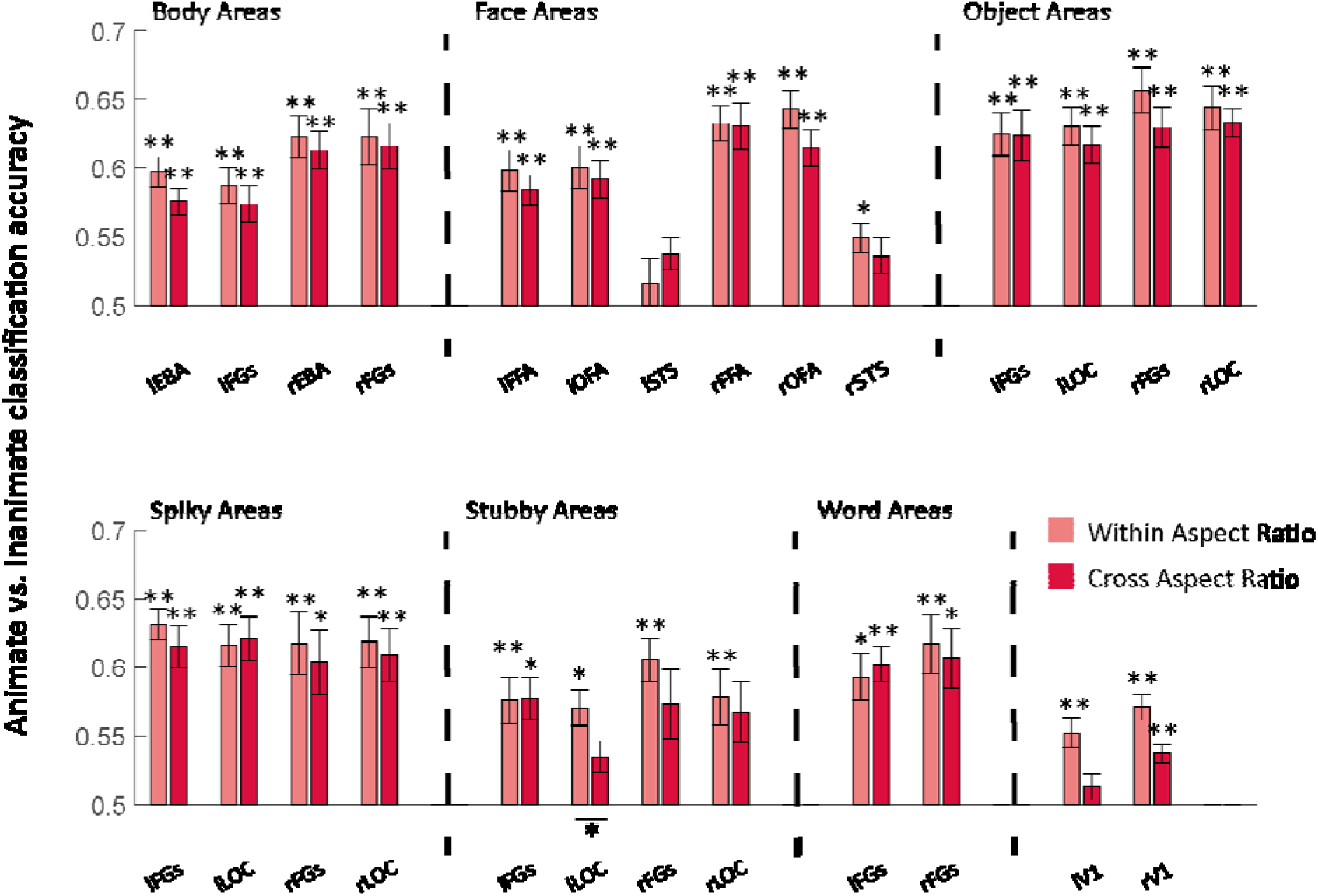
Animate vs. Inanimate classification in category-selective ROIs. The bar plot shows the mean classification accuracy for animate vs. inanimate classification when aspect ratio is similar (within) or very different (across) in the training and the test images. Error bars indicate standard errors of the means. One-sided one sample or two-sided paired t-test, FDR-corrected, * p<0.01, ** p < 0.001.]

**Figure 8.**
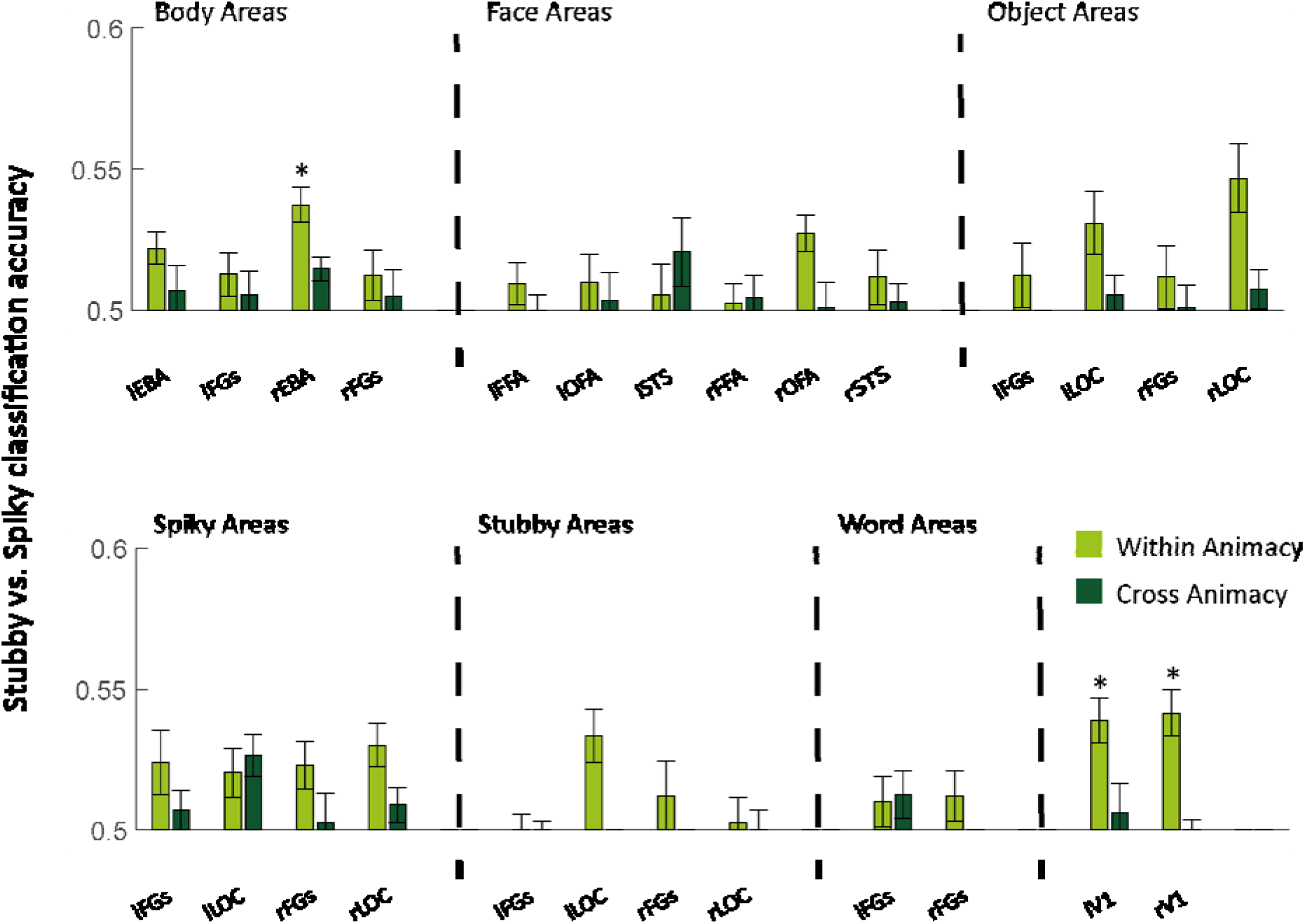
Spiky vs stubby classification in category-selective ROIs. The bar plot shows the mean classification accuracy for spiky vs stubby classification when animacy is similar (within) or very different (across) in the training and the test images. Error bars indicate standard errors of the means. One-sided one sample or two-sided paired t-test, FDR-corrected, * p<0.01.]

As could already be expected from the initial representational similarity findings, accuracies were larger for animate vs. inanimate classification than spiky vs. stubby in most OTC regions. The invariance shown in the current section in addition substantiates the claim that this salient representation of animacy stands orthogonal to aspect ratio in a strong sense of the word: It allows the determination of animacy independent from large changes in aspect ratio.

### Univariate responses reflect differences in animacy and not the aspect ratio

Category-selective regions were recognized and defined based on univariate responses and there are discussions over the dependency of these selectivities on different features. In the literature at large, it has been suggested that face-selective cortex has a preference for curved and concentric shapes^24–27^ and body-selective cortex has a preference for shapes with a comparable physical form to bodies^28,29^. More specifically based upon ref.^15^, one would expect that the face-selective cortex would prefer stubby stimuli, and the body-selective cortex would have a stronger response to spiky stimuli. With our stimulus design, we can test whether this hypothesis holds while stimulus class (e.g., face and body) and aspect ratio are properly dissociated. To evaluate the univariate effect of animacy and aspect ratio, we obtained the mean response of each category-selective ROI to each of the animacy-aspect ratio conditions (animate-spiky, animate-stubby, inanimate-spiky, and inanimate-stubby). The result (Fig 9) clearly shows that most category-selective ROIs had larger responses for animates than inanimates regardless of aspect ratio. However, comparing responses for animate-spiky with animate-stubby and responses for inanimate-spiky with inanimate-stubby, we found no significant differences in any of the category-selective ROIs such as face- and body-selective ROIs. This refutes the idea that somehow the category selectivity of these regions is related to aspect ratio and is a difficult finding for arguments that the region would have developed in that cortical location because of the existence of an aspect ratio map early in development.

**Figure 9.**
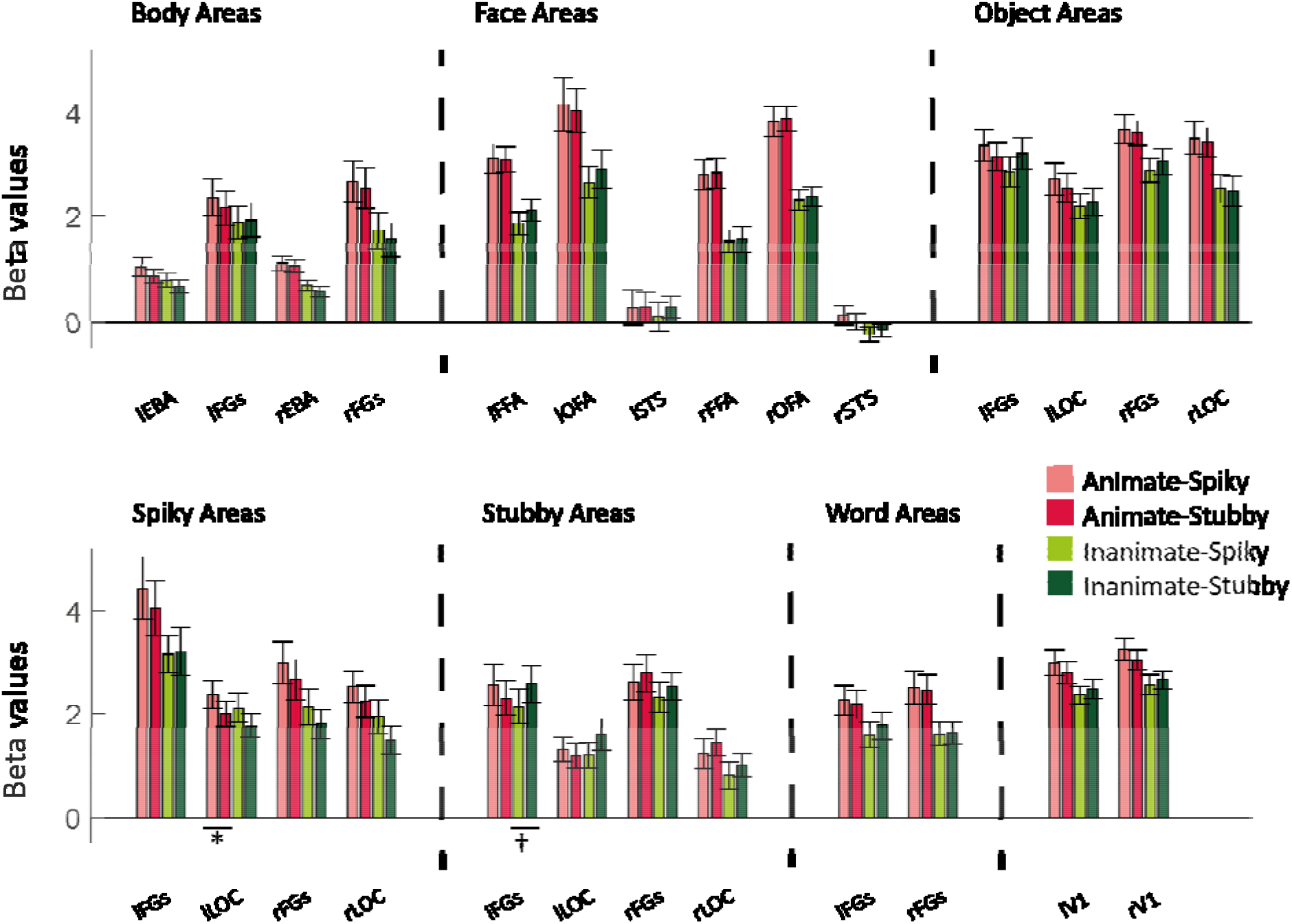
Univariate responses of category-selective ROIs to combinations of animacy and aspect ratio. Bar plots show mean beta values for animate-spiky, animate-stubby, inanimate-spiky, and inanimate-stubby. Error bars indicate standard errors of the means. Two-sided paired t-test, FDR-corrected, * p < 0.01, † p < 0.05.]

Nevertheless, there are some small regions in occipital and occipitotemporal cortex with a replicable preference for spiky or stubby objects. Several of our ROIs were selected to have such preference based on the localizer data, and some of these preferences partially replicated with our experimental stimuli. Specifically, lLOC-spiky showed a higher response for animate-spiky than for animate-stubby (Fig 9, two-sided paired t-test, FDR-corrected, t(14) = 6.55, * p < 0.01), and lFGs-stubby for inanimate-stubby !!!!! compared to inanimate-spiky (Fig 9, two-sided paired t-test, FDR-corrected across 52 tests, t(14) = 6.54, † p < 0.05). Nevertheless, this is only a partial replication of the selectivity seen in the localizer data, which already hints at how weak this selectivity is. Furthermore, while aspect ratio tuning exists in some small sub-regions of the ventral visual pathway, it falls short from explaining the large-scale organization of object representations and in particular it fails to capture the selectivity of face- and body-selective regions.

### Animacy and not the aspect ratio model explains the latent space of BigBiGAN

BigBiGAN has shown great success in generating high-quality and visually plausible images. To test if animacy and aspect ratio had key roles in determining object identity for BigBiGAN, we obtained the mapping of our stimuli to its latent space and applied PCA. Figure 10A shows the plot of the first two PCs computed for latent vectors. The separation of animate from inanimate stimuli is clearly seen, but no similar arrangement based on aspect ratio is perceivable. Representational similarity analysis quantifies this observation; the BigBiGAN RDM (Figure 10B, see method) was significantly correlated with the animacy model (r = 0.6096, one-sided permutation test p < 0.001) and not the aspect ratio model (r = 0.003, one-sided permutation test p = 0.418). Using the full pattern of representational similarity (without the data reduction step of PCA), RSA resulted in significant correlations between BigBiGAN representational similarity and both the animacy model (r = 0.3658, one-sided permutation test p < 0.001) or aspect ratio model (r = 0.1019, one-sided permutation test p < 0.01), but the correlation with the animacy model was almost 3.5 times larger than the correlation with the aspect ratio model. These data showed that animacy was the dominant factor in shaping the latent space of BigBiGAN.

**Figure 10.**
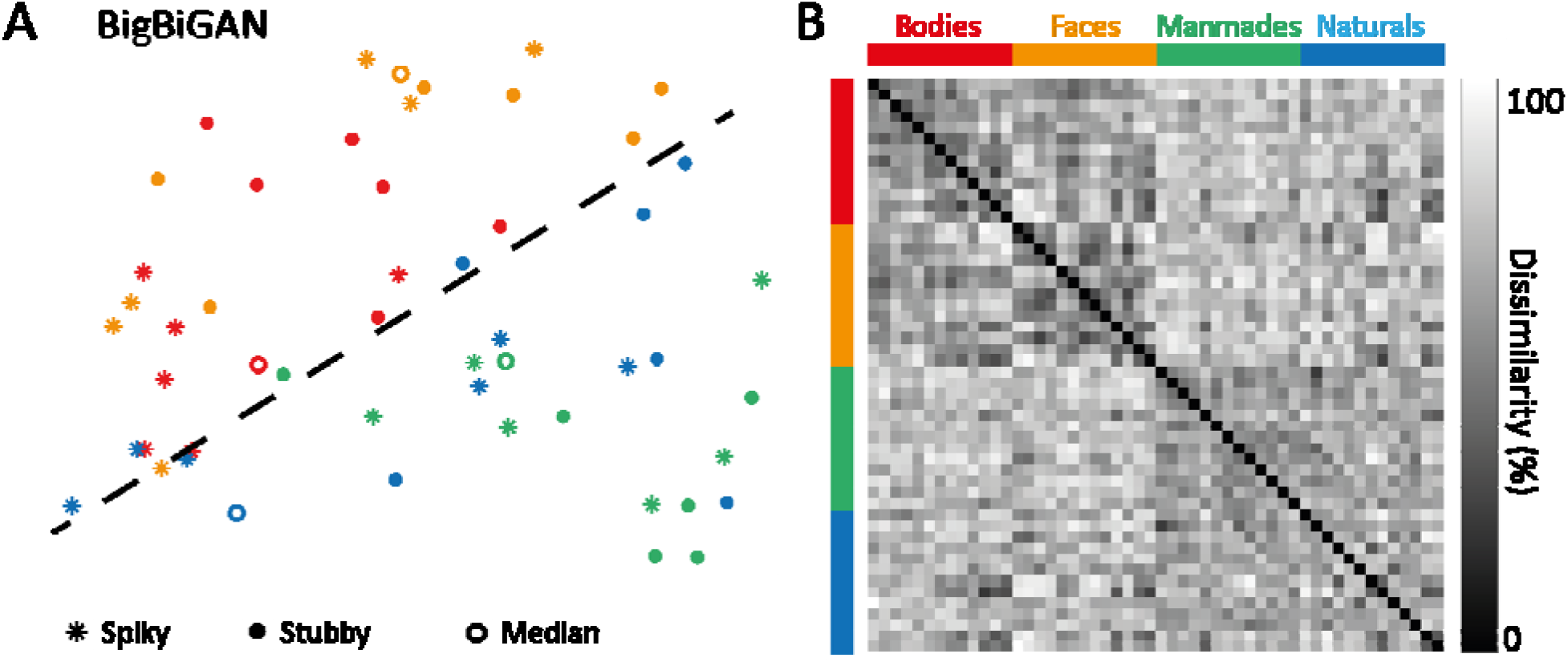
BigBiGAN’s latent vectors for our stimulus set. **A**. the first two principal components computed for the latent vectors. Points are color-coded based on stimulus category and are dots, rings, or asterisks based on aspect ratio. The dashed line shows the separation of animate from inanimate. **B**. BigBiGAN RDM. The axes of RDMs are color-coded based on stimulus category.]

We considered the possibility that we fail to find a convincing representation of aspect ratio because our stimulus set would include a too restricted range of aspect ratios. This is unlikely because of the aforementioned replication of an effect of aspect ratio specifically for inanimate objects in the part of cortex that does not preferentially represent faces and bodies. Nevertheless, fact remains that due to the need to control the aspect ratio between stimulus categories the range of aspect ratio ranges from ∼1 to ∼11 in our stimulus set. In comparison, the range was higher in the set of ref.^15^, ∼1 to ∼15, mostly due to a few outlier shapes with high aspect ratios. So, to be absolutely sure, we performed further analyses on how BigBiGAN’s representation relates to the range of included aspect ratios. We performed the equation of aspect ratio by reducing the range in the ref.^15^ stimulus set to our range (here referred to as Bao-restricted). PCA plots (supplemental figure S1) showed clear clustering according to aspect ratio for Bao-restricted. We conclude that the main experimental design property that determines the occurrence of aspect ratio clustering in BigBiGAN is whether the stimulus set confounds the distinction of faces and bodies with aspect ratio, and not the included range of aspect ratios.

## Discussion

We designed a novel stimulus set that dissociates object category (face, body, manmade and natural) from aspect ratio, and recorded fMRI responses in human OTC and DNN representations for this stimulus set to examine two alternative hypotheses: An object space organized with two main dimensions of animacy and aspect ratio, or an object space with a primary dimension of animacy and a further distinction between faces and bodies that is NOT related to the aspect ratio of these categories. Using fMRI representational similarity analysis, results show that the whole OTC and most of OTC’s category-selective ROIs represent object animacy and not aspect ratio. Restricting representational similarity analysis based on selectivity for one category reveals that object-selective ROIs in the right hemisphere slightly care about the aspect ratio of objects (manmade and natural). The represented animacy content is not constant but increases and then decreases along the anatomical posterior-to-anterior axis in VOTC. This dominant effect of animacy is reflected in larger classification accuracies for animate vs. inanimate compared to stubby vs. spiky. The animate vs. inanimate classifiers also show invariance to the aspect ratio that could express the independent representation of animacy in human OTC. Consistent with MVPA results, univariate responses of category-selective ROIs show an obvious effect of animacy but they are not influenced by aspect ratio. Similar to human OTC, BigBiGAN (the most advanced DNN in capturing object properties) represents animacy, as has been observed before^18,30–32^, but not aspect ratio. Finally, results of data-driven approaches clearly show clusters for face and body stimuli and separation of animate from inanimate stimuli in the representational space of OTC and BigBiGAN, but there is not any arrangement related to aspect ratio. In conclusion, these findings rule out the two-dimensional animacy x aspect ratio model for object space in human OTC while an object space with a primary dimension of animacy and a further distinction between faces and bodies could clearly explain the results.

We find that the organization of object space in human OTC is not related to aspect ratio. This is in contrast to the proposed model by ref.^15^ which considers stubby-spiky as one of the two main dimensions of object space in the IT cortex. There are two major differences between the present study and ref.^15^ that could explain the contrasting results; stimulus sets and subject species. In the stimulus set of ref.^15^, animate-inanimate and stubby-spiky were dissociated overall but it was not true between stimulus categories, particularly for faces and bodies: Face stimuli were all stubby and body stimuli were mostly spiky. Therefore, there was a major confound between aspect ratio and stimulus category. To overcome this problem, we had a comparatively wide range of aspect ratios for different categories and explicitly dissociated aspect ratio from these two categories. Our result showed no significant representation of aspect ratio in OTC suggesting that much of the evidence for aspect ratio as an overall dimension and as a dimension underlying face and body selectivity and the location of selectivity for these categories were due to this major confound between aspect ratio and stimulus category (face versus body).

The other major difference is species; results in ref.^15^ were mostly based on electrophysiology data from monkeys while we investigated fMRI data from human subjects. Aspect ratio might be more important for object representations in monkeys. This suggestion is supported by earlier work, although it was never investigated specifically. In particular, ref.^11^ compared monkey electrophysiology data and human fMRI for the same set of stimuli and reported common organization of object space in humans and monkeys. However, when looking more closely at their stimulus arrangement (Figure 2.A, ref.^11^), we can recognize an aspect ratio effect in monkey (spiky stimuli are closer to each other) that is not present in human (see also ref.^33^). Note however that ref.^33^ reported in a conference abstract to partially replicate the findings of ref.^15^ in humans, using the same stimulus set as ref.^15^. For that reason, we doubt that species is the main explanation for the divergence between our findings and the earlier work of ref.^15^.

Despite the absence of an overall dimension of aspect ratio, our findings are still in line with earlier reports of shape selectivity and even selectivity for aspect ratio. Previous studies on shape representation provided evidence of a relationship between perceived shape similarity and neural representation in object-selective ROIs, investigating both single-unit recordings in monkeys^34^ and fMRI data in humans^22,35–37^. Recent work has illustrated that many tens of dimensions contribute to the perception of the visual form^38^, and several of these dimensions are correlated with aspect ratio (which in the specialized literature is referred to as ‘compactness’ or ‘circularity’). A previous study that dissociated animacy from shape also suggested a role for aspect ratio in the representation of shape^2,37^. Nevertheless, we show that the face and body selectivity take priority when we properly dissociate these categories from aspect ratio. Tuning for aspect ratio is restricted to part of OTC, generally object-selective regions and a few small spots with univariate preference, and mostly restricted to inanimate objects. While aspect ratio is one of many dimensions by which object shape is represented, it does not have a special status for explaining the large-scale object space and how it is organized neuroanatomically in selective patches for e.g. faces and bodies. Other constraints that are extrinsic to the features of faces and bodies might be more important as organizing principles, including retinotopic biases induced by the pattern of eye fixation and connectivity with domain-specific systems in the brain (for review, see ref.^39^).

In face and body regions, we find no evidence for tuning for aspect ratio. There is a hypothesis that category selectivity for faces and bodies is partly explained by the aspect ratio of stimuli^24–29^. Following this hypothesis, stronger responses for stubby stimuli in face-selective and for spiky stimuli in body-selective regions are anticipated. Here, our stimulus set including faces and bodies with wide ranges of aspect ratios provides a touchstone for testing this hypothesis. We compared responses for animate-spiky with animate-stubby and responses for inanimate-spiky with inanimate-stubby and found no significant differences in any of the category-selective ROIs such as face- and body-selective ROIs. Probably, other stimulus features are more prominent when it comes to representing faces and bodies.

While our findings put doubt on the role of aspect ratio, we confirm a strong selectivity for animacy. We found that animacy is a determining factor in organizing object space in both human OTC and BigBiGAN, consistent with ref.^15^ and earlier reports (e.g., ref.^2,11^). This representation for animacy was prominent for a long distance along the anatomical posterior-to-anterior axis in VOTC. Following a cross-decoding approach, we found that animacy is represented independently of aspect ratio. Previous studies have disclosed that animacy and category distinctions are correlated with low/high-level visual features^40–43^, but there is enough evidence that animacy and category structure remain even when dissociated from shape or texture features^2,10,37,44,45^. The strength of animacy selectivity combined with a lack of aspect ratio tuning was very consistent across smaller ROIs with OTC. This indicates a high level of analogy in multivariate patterns across category-selective regions, as previously reported by ref.^46^ while they studied multivariate responses of category-selective regions (lateral occipitotemporal, occipitoparietal, FFA, PPA, EBA, lateral occipital) to eight different categories (bodies, buildings, cars, cats, chairs, faces, hammers, and phones).

The occipitotemporal cortex has a key role in visual object recognition, but the organization of object space in this region is still unclear. To examine hypotheses considering animacy, aspect ratio and face-body as principal dimensions characterizing object space in the occipitotemporal cortex, we devised a novel stimulus set that dissociates these dimensions. Investigation of human fMRI and DNN responses to this stimulus set shows that a two-dimensional animacy x aspect ratio model cannot explain object representations in either occipitotemporal cortex or a state-of-the-art DNN while a model in terms of an animacy dimension combined with strong selectivity for faces and bodies is more compatible with both neural and DNN representations.

## Methods

### Participants

Seventeen subjects (12 females, 19–46 years of age) took part in the experiment. All subjects were healthy and right-handed with normal visual acuity. They gave written informed consent and received payment for their participation. The experiment was approved by the Ethics Committee on the use of human subjects at the Universitair Ziekenhuis/Katholieke Universiteit Leuven. Two subjects were later excluded due to excessive head motion (see Preprocessing for more details), and the final analyses included the remaining 15 subjects.

### Stimuli

Stimuli in the main experiment included 52 images from four categories: animal body, animal face, manmade and natural object (Fig. 1A). We selected 13 exemplar images from each of these four categories that were selected to provide a comparably wide range of aspect ratios for different categories. In the literature on shape description, aspect ratio is usually defined as the ratio of principle axes in a shape, but following ref.^15^, we defined aspect ratio as a function of perimeter P and area A; aspect ratio = P^2^/(4πA). Using this formula, the ranges of aspect ratio for body and face stimuli in ref.^15^ were (1.38-6.84) and (0.91-1.19), respectively while we had ranges of (1-9.66) for bodies and (1.51-10.89) for faces. In each category, we sorted stimuli according to the aspect ratio, determined the category median on aspect ratio, and labeled stimuli with aspect ratios larger than the category median as spiky and those with aspect ratios smaller than the category median as stubby.

All images were grayscale with a gray background, cropped to 700*700 pixels (subtended 10 degrees of visual angle in the MRI scanner). We equalized the luminance histogram and the average energy at each spatial frequency across all the images using the shine toolbox^47^. Examples of finalized stimuli used for fMRI experiments and computational analyses are provided in Figure 1B.

To compare stimuli with respect to animacy, aspect ratio, and low-level shape properties, we computed representational dissimilarity matrices (RDMs) and quantified pair-wise resemblance of images for these properties. For constructing a dissimilarity matrix based on animacy, we assigned scores ‘+1’ to animate (face and body) and ‘-1’ to inanimate (manmade and natural) stimuli and computed the absolute difference of scores for each stimulus pair (Fig 1C). The aspect ratio model included the pairwise absolute difference of aspect ratios (Fig 1C). As measures of low-level shape properties, we investigated pixelwise dissimilarity^22^ and dissimilarities based on outputs of hmax model (S1-C1, C2)^48^. We computed correlation distances between pixels/S1-C1/C2 for each pair of stimuli to build dissimilarity matrices. Comparison of model RDMs (Spearman correlation and p-values from one-sided permutation tests) showed non-significant similarity between animacy and aspect ratio models (r = −0.0201, p = 0.8800), animacy and low-level models (pixels: r = 0.0358, p = 0.1150, S1-C1: r = 0.0412, p = 0.0540, C2: r = − 0.0052, p = 0.5270), and aspect-ratio and low-level models (pixels: r = −0.0057, p = 0.5460, S1-C1: r = 0.0148, p = 0.3600, C2: r = 0.0473, p = 0.2080).

### Scanning procedures

Data recording consisted of eight experimental runs, three localizer runs, and one anatomical scan, all completed in one session for each participant.

For the experimental runs, we used a rapid event-related design. Each experimental run included a random sequence of 138 trials; two repetitions of each stimulus image plus 34 fixation trials. Each trial was 3 s. Stimulus trials began with the stimulus presentation for 1500 ms and were followed by 1500 ms of the fixation point. Each experimental run lasted for 7 min and 14 s. The fixation point was presented at the center of the screen continuously throughout each run. Subjects were instructed to fixate on the fixation point and, on each trial, press a button to indicate whether they preferred looking at the current image or the previous one (same task as in ref.^49^).

For the localizer runs, we used a block design with six stimulus types: body, face, stubby object, spiky object, word, and box-scrambled version of the object images. Each run included 24 blocks with four blocks for each stimulus type. The presentation order of the stimulus types was counterbalanced across runs. Each block lasted for 16 s. There was a 10-s blank period in the beginning and end, and three 12-s blank periods between stimulus blocks of each repetition. Each localizer run lasted for 7 min and 20 s. In each stimulus block, there were 18 images of the same stimulus type and two different randomly selected images were repeated. Each image in a block appeared for 400 ms followed by 400ms fixation. A fixation point was presented at the center of the screen continuously throughout each run. Subjects were instructed to fixate on the fixation point, detect the stimulus one-back repetition, and report it by pressing a response key with their right index finger. For one subject, the data for one of three localizer runs were discarded since there were uncorrectable artifacts in the data. Subjects viewed the visual stimuli through a back-projection screen, and the tasks were presented using MATLAB and Psychtoolbox-3^50^.

### Acquisition parameters

Magnetic resonance imaging (MRI) data were collected at the Department of Radiology of the Universitair Ziekenhuis Leuven university Hospitals using a 3T Philips scanner, with a 32-channel head coil. Functional images were obtained using a two-dimensional (2D) multiband (MB) T2*-weighted echo planar imaging sequence with an MB of 2, time repetition (TR) of 2 s, time echo (TE) 30 ms, 90⍰ flip angle, 46 transverse slices, and a voxel size of 2×2×2 mm^3^. A high-resolution T1-weighted structural scan was also acquired from each participant using an MPRAGE pulse sequence (1×1×1 mm^3^ isotropic voxels).

### Preprocessing

fMRIPrep^51^ was employed for preprocessing anatomical and functional data, using default settings unless otherwise noted. The T1-weighted image was corrected for intensity non-uniformity, skull-stripped, and went through nonlinear volume-based registration to ICBM 152 nonlinear Asymetrical template version 2009c. Each of the bold runs was motion-corrected, coregistered to the individual’s anatomic scan, and normalized into standard space MNI152NLin2009cAsym. Subjects with excessive head motion (frame-wise displacement>2 mm; more than one voxel size) were excluded (two subjects). Subjects either did not have frame-wise displacements greater than the predefined threshold or had it repeated multiple times. The rest of the analyses were conducted with SPM12 software (version 6906). As the last step of preprocessing, all functional volumes were smoothed using a Gaussian kernel, 4 mm FWHM. After preprocessing, a run-wise general linear model (GLM) analysis was performed to obtain the beta values for each stimulus image of the experimental and each stimulus type of the localizer runs in each voxel. For the experimental runs, the GLM consisted of the 52 stimulus regressors (boxcar functions at the stimulus onsets with a duration of 1500 ms convolved with a canonical hemodynamic response function) and six motion correction parameters (translation and rotation along the x-, y-, and z-axes). For the localizer runs the GLM included the six stimulus regressors (boxcar function at the block onsets with a duration of 16 s convolved with a canonical hemodynamic response function) and the same six motion correction parameters.

### Defining regions of interest (ROIs)

We used contrasts from localizer runs intersected with masks from functional^52^ or anatomical atlases (Anatomy Toolbox^53^) to define a maximum of 26 ROIs covering lateral and ventral OTC in each individual subject. Since the functional atlas included few ROIs in the ventral cortex, we used the intersection of contrast maps with FG1-FG4 (FGs) from the Anatomy Toolbox to define more ventral ROIs. All lateral and ventral body, face, and object selective ROIs in each hemisphere were merged together to produce a large ROI representing left or right OTC. We also used the anatomical V1 mask in defining EVC. Details on localizing ROIs (employed functional contrasts and masks) are provided in supplemental table S1. To ensure that ROIs with the same selectivity were independent, we examined their overlap and took overlapping voxels away from the larger ROIs. This mostly happened for LOC and FGs in the object, spiky, and stubby areas and overlapping voxels were removed from LOC. Note that this removal of overlap did not meaningfully change any of the findings and statistics, and very similar findings were obtained when these voxels are not removed.

To capture gradual effects, we defined a series of 37 small consecutive ROIs along the antero-posterior axis that forms the ventral visual stream. These ROIs were localized using the functional contrast of ‘All conditions>Fixation’ intersected with anatomical masks including V1, V2, V3v, V4v and FGs^54^.

All ROIs included at least 25 voxels that surpassed the statistical uncorrected threshold *p* < 0.001 in the relevant functional contrast and were included in the relevant mask (supplemental table S1). If the number of survived voxels was less than 25, a more liberal threshold of *p* < 0.01 or *p* < 0.05 was applied.

Figure 3 and Figure 6 respectively show category-selective ROIs and the ventral visual stream in one representative subject mapped onto the inflated cortex using FreeSurfer (https://surfer.nmr.mgh. harvard.edu).

Mean sizes (standard errors) for category-selective ROIs were as follows: lEBA = 823(54), rEBA = 1064(77), lFGs-body = 145(28), rFGs-body = 138(26), lFFA = 90(13), rFFA = 139(19), lOFA = 54(7), rOFA = 122(13), lSTS = 39(7), rSTS = 91(29), lFGs-object = 326(50), rFGs-object = 208(31), lLOC-object = 294(42), rLOC-object = 284(44), lFGs-spiky = 99(28), rFGs-spiky = 77(16), lLOC-spiky = 248(61), rLOC-spiky = 207(74), lFGs-stubby = 50(8), rFGs-stubby = 53(6), lLOC-stubby = 83(31), rLOC-stubby = 70(9), lFGs-word = 207(38), rFGs-word = 50(6), lV1 = 335(24), rV1 = 467(22).

### Statistical analysis of main fMRI experiment

Multivariate (representational similarity analysis and classification) and univariate analyses were used to study the role of animacy and aspect ratio in organizing OTC’s object space. In-house MATLAB code and CoSMoMVPA^55^ toolbox were employed for the following analyses.

#### Representational similarity analysis

We compared the representational dissimilarity matrices (RDMs) based on neural activity in different ROIs with model RDMs based on animacy and aspect ratio. Neural RDM for each ROI included the pairwise Mahalanobis distance between activity patterns (beta weights) of the ROI for different stimuli^56,57^. The off-diagonal of the neural and model RDMs were vectorized and the Spearman’s correlation between dissimilarity vectors was then calculated and compared. We tested the significance of correlation values between neural and model RDMs across subjects employing one-sample t-tests and the significance of differences between correlation values obtained for animacy RDM and aspect ratio RDM across subjects using paired t-tests. Then, p-values were corrected for multiple comparisons across all ROIs (false discovery rate^58^).

For body, face, and object selective ROIs, we repeated the representational similarity analysis for a subset of our stimuli limited to their favorite category; neural and aspect ratio RDMs were computed using body, face or object (manmade and natural) stimuli separately, the off-diagonal of the neural and model RDMs were vectorized, the Spearman’s correlation between dissimilarity vectors was calculated and tested for significancy employing one-sample t-tests using correction for multiple comparisons across ROIs (false discovery rate^58^).

When correlating neural and model dissimilarity, we can compare this correlation with the reliability of the neural dissimilarity matrix. This reliability can be interpreted as the estimate of the highest possible correlation given the noise in the data^59^. For each region, it was computed as the cross-validated correlation of each subject’s neural dissimilarity matrix with the mean of the remaining subjects’ neural dissimilarity matrices. These reliabilities are provided as gray background bars in all the relevant figures.

For OTC, we averaged the neural dissimilarity matrices across subjects to obtain a group-level neural dissimilarity matrix, applied multidimensional scaling (MDS) analysis, and used the first two dimensions that explained most of the variance to produce an MDS diagram in which the distance among stimuli expresses the similarity among their neural representation.

#### Within and cross-category classification

We performed category classification i.e. animate vs. inanimate and spiky vs. stubby classification. In a leave-one-run-out cross-validation procedure, samples (beta weights) were partitioned to train and test sets and linear discriminant analysis (LDA) was employed to perform brain decoding classification in individual brains. The animate vs. inanimate classification was performed within and cross aspect ratio; for within aspect ratio, we trained and tested classifiers with the same category of stimuli (either spiky or stubby) while for cross aspect ratio, classifiers were trained and tested with stimuli from different categories (supplemental table S2). The spiky vs. stubby classification was also performed within and cross animacy; for within animacy, we trained and tested classifiers with the same category of stimuli (either animate or inanimate) while for cross animacy, classifiers were trained and tested with stimuli from different categories (supplemental table S2). We tested if the classification accuracies for within or cross-category classifications across subjects were significantly above chance level (50% chance level for two-category classification) employing one-sample t-tests and the significance of differences between within and cross-category classification accuracies across subjects using paired t-tests. Then, p-values were corrected for multiple comparisons across ROIs (false discovery rate^58^). Comparing within and cross-category classification accuracies for these two dimensions, we determined ROIs’ capability of generalizing distinction along one dimension over the other dimension / specified ROIs in which animacy and aspect ratio contents are independently represented.

#### Univariate analysis

We investigated the mean response of each category-selective ROI to each of the animacy-aspect ratio conditions; animate-spiky, animate-stubby, inanimate-spiky, and inanimate-stubby. For each stimulus, beta weights were averaged across all runs, and then all voxels within each ROI in individual subjects. Then, responses for animate-spiky were compared with animate-stubby and responses for inanimate-spiky were compared with inanimate-stubby employing paired t-tests across participants, and p-values were corrected for multiple comparisons across ROIs (false discovery rate^58^).

#### Computational simulation with artificial neural network: BigBiGAN

BigBiGAN is the state-of-the-art artificial neural network for unconditional image generation regarding image quality and visual plausibility. It is a large-scale bi-directional generative adversarial network, pre-trained on ImageNet that converts images into a 120-dimensional latent space. This unified latent space captures all properties of objects including high-level image attributes and categories^60,61^. In order to study features that characterize the identity of our stimuli for BigBiGAN, we passed our stimuli through the network (https://tfhub.dev/deepmind/bigbigan-revnet50X4/1) and acquired corresponding latent vectors. Then, for obtaining the main factors, principal component analysis (PCA) was applied to the latent vectors and the first two principal components (PCs) were used for visualization. We computed an RDM based on correlation distances of the first two PCs (called BigBiGAN RDM), vectorized its off-diagonal and calculated Spearman’s correlation between this dissimilarity vector and the vectors from the animacy and aspect ratio model. The significance of these correlations is also verified by employing permutation tests.

## Acknowledgments

This work was supported by: Fonds voor Wetenschappelijk Onderzoek (FWO, 12A6122N, G0D3322N, G073122N), Excellence Of Science (EOS grant G0E8718N). The authors thank A. Platonov for help with setting up the study.

## Author contributions

E.Y and H.O.B. developed the study design and analysis plan, E.Y. collected and analyzed data, and E.Y and H.O.B. wrote the manuscript.

## Competing interests

The authors declare no competing interests.

## Supplementary Information

## Supplementary Tables

**Supplementary table S1.** Functional contrasts and masks used to localize category-selective ROIs.

| ROIs | Contrast | Mask |
| --- | --- | --- |
| Body areas |  |  |
| IFGs-body & rFGs-body | body>(face+objects<sup>a</sup>) | anatomical atlas |
| IEBA & rEBA |  | functional atlas |
| Face areas |  |  |
| IFFA & rFFA | face>(body+object<sup>a</sup>) | functional atlas |
| IOFA & rOFA |  | functional atlas |
| ISTS & rSTS |  | functional atlas |
| Object areas |  |  |
| IFGs-object & rFGs-object | object<sup>a</sup>>scramble | anatomical atlas |
| lLOC-object & rLOC-object |  | functional atlas |
| Spiky areas |  |  |
| IFGs-spiky & rFGs-spiky | spiky>stubby | anatomical atlas |
| lLOC-spiky & rLOC-spiky |  | functional atlas |
| Stubby areas |  |  |
| IFGs-stubby & rFGs-stubby | stubby>spiky | anatomical atlas |
| lLOC-stubby & rLOC-stubby |  | functional atlas |
| Word areas |  |  |
| IFGs-word & rFGs-word | word>object<sup>a</sup> | anatomical atlas |
| EVC |  |  |
| lV1 & rV1 | All conditions>Fixation | anatomical atlas |

<sup>a</sup>Object condition includes both spiky and stubby objects.

**Supplementary table S2.** Train and test sets for within and cross-category classification.

| Classifier |  | Train set | Test set |
| --- | --- | --- | --- |
| Animate-inanimate | within aspect ratio | spiky | spiky |
|  |  | stubby | stubby |
|  | cross aspect ratio | spiky | stubby |
|  |  | stubby | spiky |
| Spiky-stubby | within animacy | animate | animate |
|  |  | inanimate | inanimate |
|  | cross animacy | animate | inanimate |
|  |  | inanimate | animate |

## Supplementary Figures

**Supplemental Figure S1.**
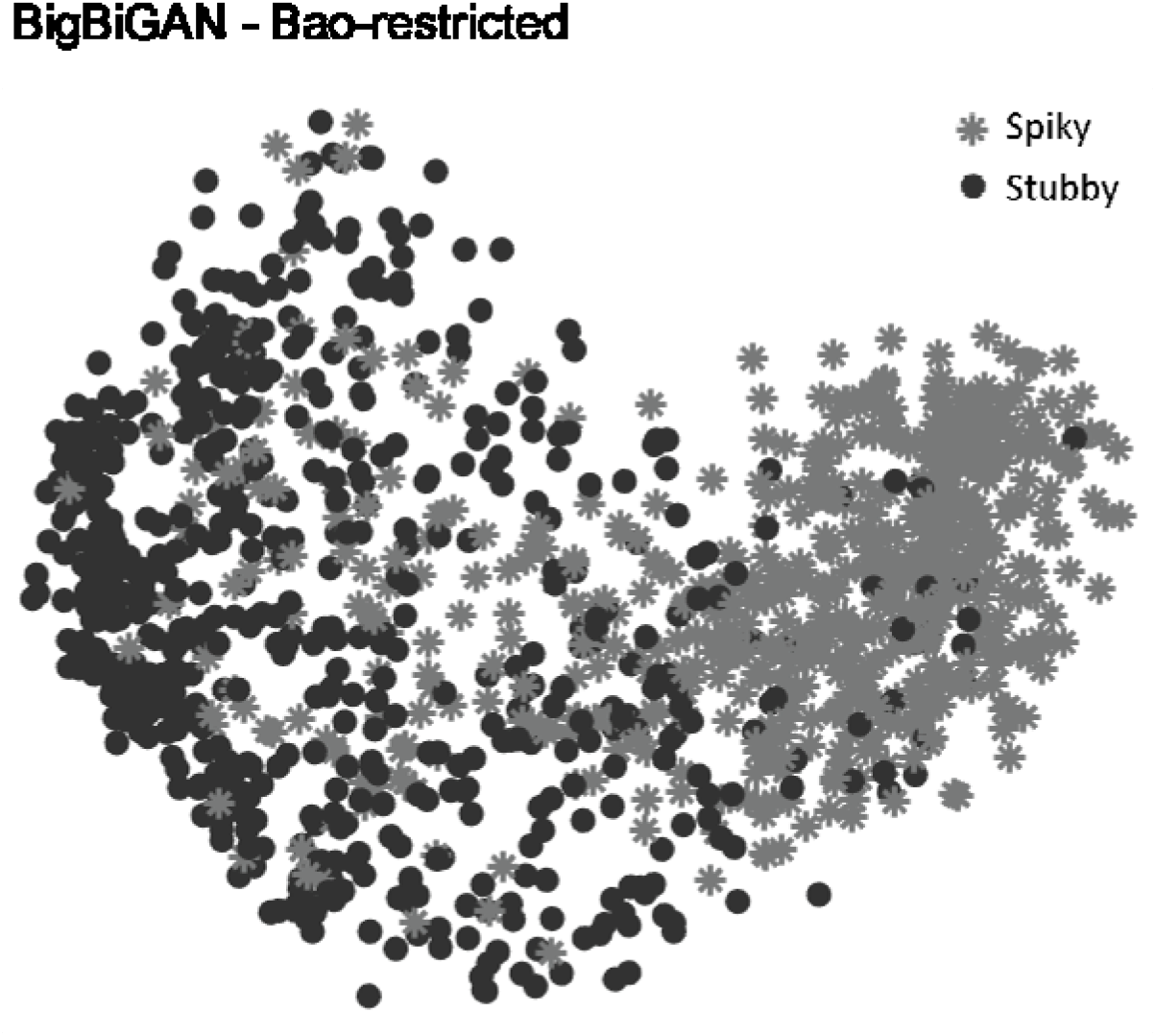
BigBiGAN’s latent vectors for Bao-restricted stimulus set. The first two principal components computed for the latent vectors. Points are color/shape-coded based on aspect ratio.

## Notes

### Competing Interest Statement

The authors have declared no competing interest.

